# Pretrained protein language models choose between sequence novelty and structural completeness

**DOI:** 10.1101/2025.10.01.679905

**Authors:** Arjuna M. Subramanian, Zachary A. Martinez, Matt Thomson

**Affiliations:** Division of Biology and Biological Engineering, California Institute of Technology, CA, USA

## Abstract

Protein language models (PLMs) have gained increasing acceptance in tasks ranging from variant effect prediction in disease to optimization and *de novo* design of proteins with improved stability, target-binding affinity, and catalytic performance. Despite encouraging performance in such applications, little is understood as far as the degree to which PLM-generated sequences – putative novel protein outputs – recapitulate the broad biophysical rules and diversity of sequence, structure, and function that defines natural protein-space, vital knowledge for boosting the design capacity of PLMs in ever-more-complex systems. Towards this end, we computationally profile and characterize the sequence and structure statistics and properties of hundreds of thousands of potential small proteins proposed through free unconstrained generation from architecturally distinct PLMs. We show that although these models exhibit a prodigious latent capacity to access novel amino-acid sequences, they struggle to approach the structural variation that exists on plain display in nature. Moreover, we uncover a stark tradeoff between prioritizing sequence novelty or structural breadth, exemplified by a “helical bundle trap” that dominates model output when aiming outside the comfortable bounds and evolutionary organization of natural sequences. These findings underscore a critical need for strategies that can rapidly guide PLMs into unlocking through generation the full richness of protein sequence, structure, and function that is consistent with governing biophysics but tantalizingly untapped as of yet in design contexts.

**Author summary:** Large language models (LLMs) like GPT aren’t just for human text. Spinoff versions that treat protein and DNA sequences as special “languages” of their own, complete with preferred words and grammars, are being used to identify disease-causing genes and mutations and to design new treatments and drugs for clinical testing. But as anyone who has used an LLM chatbot has probably experienced at one time or another, these models can act narrow-minded or nonsensical, reducing their generality and utility. Protein language models are no exception to such flaws of reasoning. We show that protein language models face a fundamental choice between suggesting novel sequences that look nothing like natural ones and capturing the full range of three-dimensional shapes and structures responsible for the diverse functioning of molecular machines. Managing or bypassing this tradeoff is consequently of high import for designing proteins that impart novel functions and activities for therapeutic targeting and beyond.

## Introduction

Protein language models (PLMs) and genomic language models (GLMs) have become increasingly utilized as descriptors and generators of real and synthetic biological components [1–5]. Treating proteins as amino-acid “texts,” PLMs internalize knowledge about sequence determinants within natural proteins, and implicitly learn features of structure and function including *α*-helical vs *β*-strand content, residue-wise contact maps, and binding/active sites [6–8]. At the same time, they retain an innate exploratory capacity to reach novel protein structures and functional variants [2, 9–12]. This tension between exploitation and exploration points to PLMs as natural vehicles for answering fundamental questions of order and organization in the protein sequence-structure map and for engineering new proteins with improved and/or tunable properties from thermostability to catalytic turnover. It remains poorly understood, however, to what degree PLM generative outputs are consistent with natural protein-space as far as sequence statistics, structure statistics, and biochemical and biophysical properties, all essential factors in deciding how best to leverage PLMs for designing novel proteins that are plausible, useful, and functional.

The ability of PLMs to faithfully and representatively capture such statistics, stretching across the levels of sequence, structure, and function, is generally assumed to follow from two factors. First, the depth and volume of large training datasets such as the ≈ 50/90/100 million sequences in UniRef50/90/100 [13]. And second, the retention and application of transformer blocks – originally developed for large language model (LLM) work on human-created text – to detect higher-order correlations, patterns, and determinants across sequence positions [14–16]. However, this faith looks past the fact that training approaches and hyperparameters are often reused directly from LLMs without adjusting for any imperfections in the analogy between human and protein languages. Model sizes (10^7^ − 10^11^ parameters) and architectures (i.e. autoencoder vs encoder-only vs decoder-only), alphabet/vocabulary discretization (e.g. decomposing sequences into individual amino-acids or longer subsequences), and training task (i.e. autoregressive vs masked prediction objectives) vary significantly [1–3, 10, 17, 18]. So do sequence sampling and generation methods, from probabilistic left-to-right construction appending tokens conditioned on whatever has come before, to Gibbs sampling from position-wise amino-acid likelihoods in arbitrary order, to indirect schemes incorporating PLM-derived metrics into Markov Chain Monte Carlo (MCMC) searches on traditional sequence landscapes, to direct conversion of high-dimensional latent space representations back to full-length amino-acid sequences in suitable cases. [2, 10–12, 19–21]. PLMs may indeed be learning and storing energy functions and co-evolutionary statistics deep within stacked and layered transformers [22, 23]. But does that understanding manifest in the boundless synthetic protein “texts” that can now be generated at the push of a button?

To address this gap, we perform an at-scale *in silico* statistical characterization of sequence and structure composition in PLM-generated amino-acid sequences. We demonstrate that not all models and sampling strategies are created equal. In particular, autoregressive generation from a decoder-based model (ProtGPT2) outperforms Gibbs sampling from an encoder-based model (ESM2) at proposing realistic protein structures and achieving structural diversity. Relative success aside, however, the structural coverage of ProtGPT2 still sharply distorts the distribution of natural protein structures. Further, we discover that while ProtGPT2 displays an impressive ability to sample and assemble novel sequence motifs, maximizing sequence novelty through hyperparameter tuning exacerbates its already substantial shortcomings as far as preserving structural breadth, revealing a considerable tradeoff between PLM-driven diversification of sequence vs structure. Our findings highlight a critical need for PLM-based generative strategies that accurately capture rare, malleable, and novel protein features if we are to push the boundaries of fundamental biophysics and protein design as a field.

## Results

### Creating a fold-annotated database of PLM-generated protein structures

In order to characterize sequence and structure statistics, we initially constructed a database of AI-generated artificial protein sequences from a suite of representative models. We selected two commonly-used PLMs, both transformer-based but otherwise starkly contrasting in architecture and compatible sequence generation methods: (1) ProtGPT2, an autoregressive decoder-only model with 774M parameters; and (2) ESM2, a bidirectional encoder-only model that includes 8M, 35M, 150M, 650M, 3B, and 15B parameter versions, working with the 150M parameter version specifically to manage compute time overhead for sequence generation (Fig. 1) [1, 2]. Sequence generation from ProtGPT2 involves stepwise addition of tokens drawn probabilistically from a SwissProt-extracted vocabulary of 50,256 short amino-acid subsequences, proceeding left-to-right while conditioning on the in-progress sequence to the left. We sampled 100,000 sequences with ProtGPT2, applying the default best-performing hyperparameters from the original study, enforcing a target length of 100aa, and removing sequences containing rare or ambiguous amino acids (refer to Methods for full sampling/hyperparameter details).

**Fig 1.**
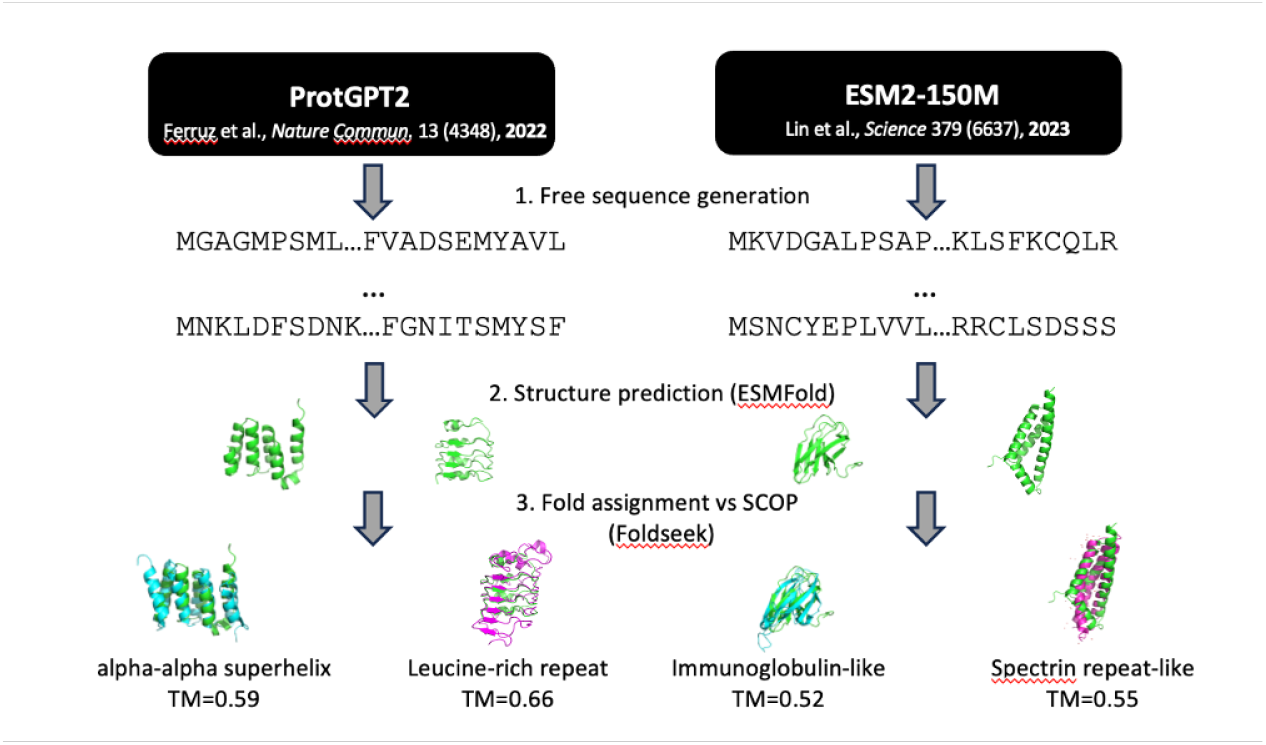
Workflow for structural annotation of pLM-generated sequences. Schematic overview of pLM generation, structure prediction, and structural search+assignment pipeline for representative models ProtGPT2 and ESM2-150M.

For ESM2-150M, we elected a left-to-right Gibbs sampling approach in single-token increments for ease of fair comparison to the autoregressive method and to align with existing benchmarks in the field [19]. In contrast to ProtGPT2 sampling, ESM2-150M uses the amino-acid alphabet as its vocabulary and generates up to a fixed sequence length. We generated 148,500 sequences of target length 100aa from ESM2-150M, with default hyperparameters and the same filtering for rare/ambiguous amino acids as before (refer to Methods for full sampling/hyperparameter details).

We also generated a control set of 74,250 random amino-acid sequences of fixed length 100aa, weighting sampling probability proportionally to natural amino-acid abundances in SwissProt, i.e. preserving first-order sequence statistics but none of the second- and higher-order correlations between residues that transformers are expected to capture. Lastly, to benchmark against structure-conditioned sequence generation, we added an inverse-folding comparison set comprised of 110,700 sequences, three per each of the 36,900 representative experimental structures in the **S**tructural **C**lassification **o**f **P**roteins (SCOP) database, designed via backbone-to-sequence inference by ESM-IF1 following default hyperparameters (refer to Methods for full sampling/hyperparameter details) [24, 25].

We predicted structures for all ∼ 430, 000 sequences with ESMFold. ESMFold has been shown to exceed the prediction accuracy of AlphaFold2 in the absence of deep multiple sequence alignment (MSA) information, which far-from-natural sequences lack by definition [1, 26]. ESMFold’s MSA-less single-sequence transformer architecture is additionally suitable for efficient inference in large-scale structure prediction tasks and for more transparent analysis of model behavior. We further assessed the ability of ESMFold to generalize to sequences outside of the natural distribution by evaluating model prediction accuracy on *de novo* designed proteins with structures deposited in the Protein Data Bank (PDB) posterior to the ESMFold training horizon of 05-01-2020.

This validation set included *n* = 122 proteins, the products of phylogenetic, physics-based, and generative AI models, with structural diversity distributed across all-*α*, all-*β* and mixed-*αβ* topologies. Alignment comparison of ESMFold predicted structures and ground-truth PDB experimental structures attained a median backbone root-mean-square deviation (RMSD) of 0.92 ± 0.14 Å, a strong indication that ESMFold sufficiently generalizes beyond natural training data to serve as a structure prediction oracle on PLM-generated sequences (Fig. 2).

**Fig 2.**
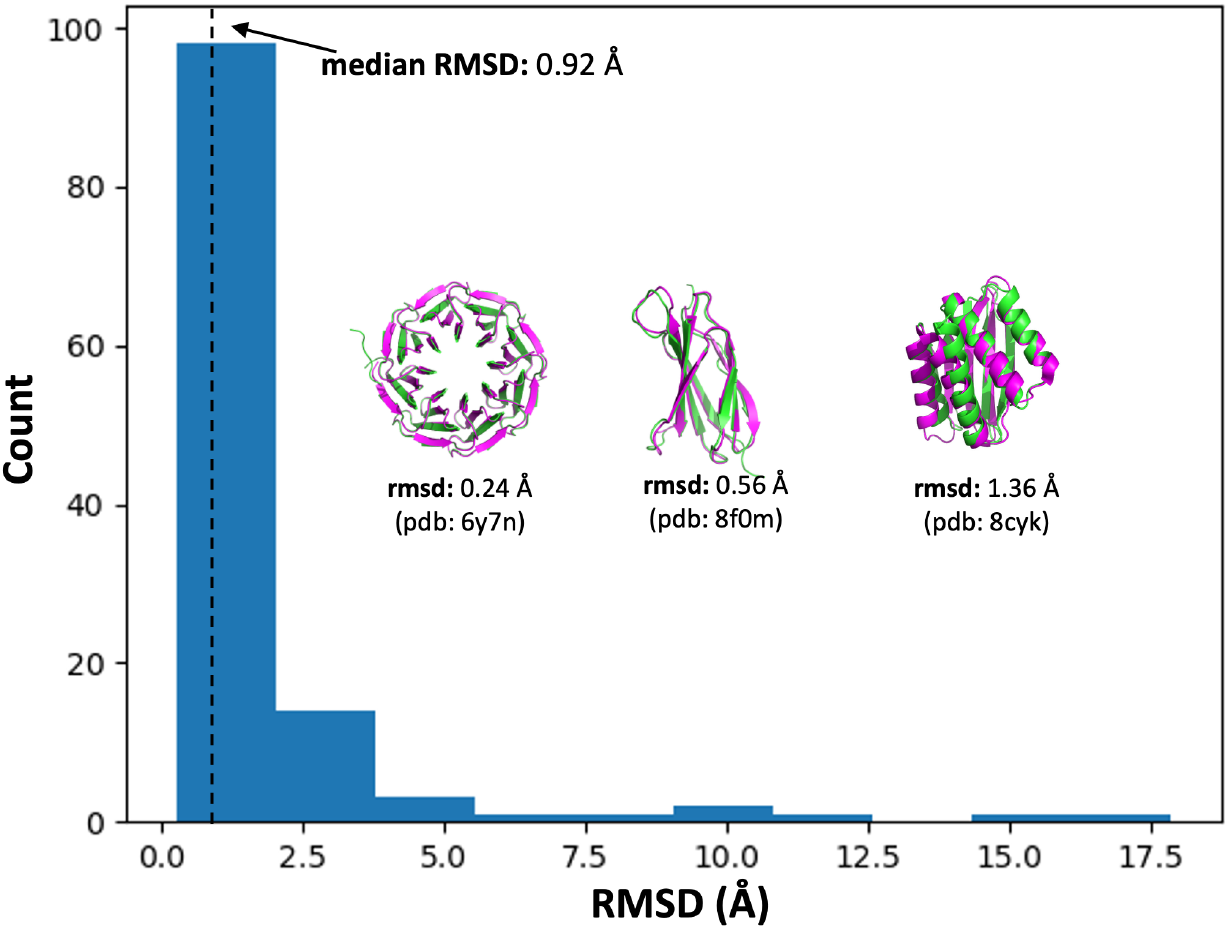
ESMFold achieves single-angstrom structure prediction accuracy on *de novo* designed sequences. Backbone atom root-mean-square deviation (RMSD; median= 0.92 ±0.14Å) for ESMFold predicted structures of *n* = 122 *de novo* designed proteins vs. experimental ground-truth structures, covering *α, β*, and mixed-*αβ* global topologies, and including designs obtained from physics-based and generative AI models. All sequences in the validation set had experimental structures deposited in the Protein Data Bank (PDB) *after* the ESMFold training cutoff date of 05-01-2020.

Finally, to identify common natural structural motifs in PLM-generated, inverse-folded, and control variants, we annotated our database of predicted structures at the “fold” level of the SCOP classification, covering 1579 possible fold labels. Each predicted structure was assigned to a consensus fold label wherever possible by performing a structure-based search against all SCOP superfamily representative PDB structures (*n* = 36, 900; the same structures used as backbone templates for inverse-folding) with Foldseek [27]. The full generation, folding, and annotation workflow is summarized in Fig 1.

### PLM-generated sequences are protein-like

For a first-pass analysis, we consider whether generated sequences and their corresponding (predicted) structures recapitulate the global characteristics of natural proteins. To assess where pLM-generated sequences lie with respect to natural sequence-space, we extract the ESM2-150M final hidden-layer internal representations (“embeddings”) of all > 400, 000 generated sequences and 100, 000 diverse natural sequences coding for SCOP fold examples mined from the AlphaFoldDB [28]. We reduce dimensionality to 2D using UMAP, and apply a rule-of-thumb that the embeddings of qualitatively similar sequences should co-localize [29]. We observe that ProtGPT2-generated sequences separate into two subpopulations, one co-localizing with natural sequences, and a second co-localizing with random sequences (Fig. 3A). ESM2-150M-generated sequences, in contrast, co-localize substantially with random sequences. Inverse-folded sequences from ESM-IF1 largely mirror the distribution of natural sequences, implying that they do not represent any significant departure from natural protein-space. This is not wholly unexpected given the tight constraints imposed by backbone-conditioning and evidence that crystallographic refinement may impart a memory of natural sidechain identities; it underscores the nature of PLM exploratory capacity as enabled by free generation without explicit constraints or conditioning in inference [30].

**Fig 3.**
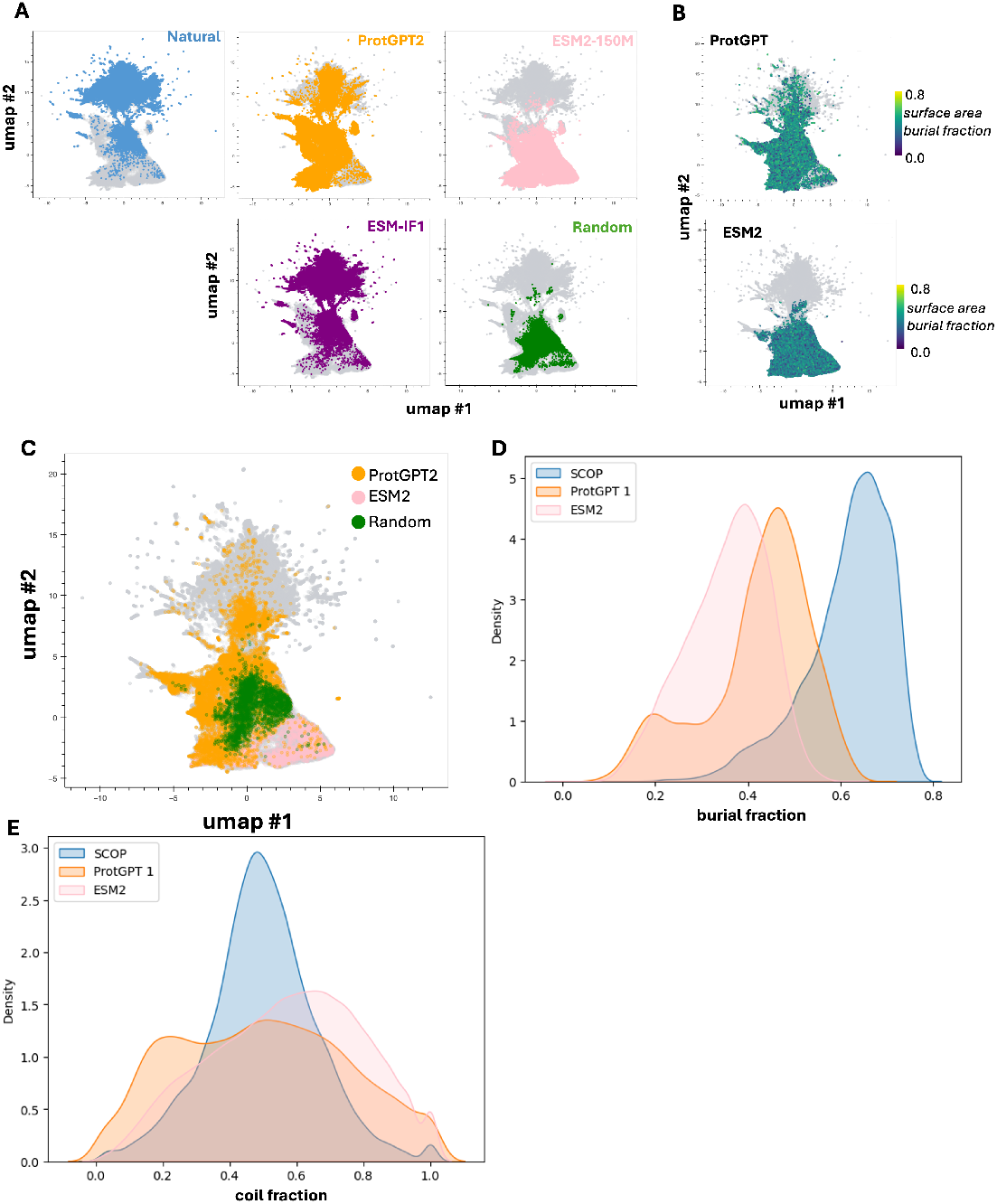
PLM-generated sequences reflect the basic properties of compact, globular proteins. (**A**) Dimension-reduced UMAP representation of ESM2-150M embeddings of natural, pLM-generated, inverse-folded, and random control sequences. (**B**) UMAP representation of pLM-generated sequences, colored by the fraction of amino-acid surface area buried (a measure of protein compactness). (**C**) UMAP representation of pLM-generated and random sequences assignable to a SCOP fold. (**D**) Fraction of amino-acid surface area buried for natural and pLM-generated sequences. (**E**) Fraction of residues annotated as random coils by DSSP for natural and pLM-generated sequences.

Turning towards coarse structural properties, the compactness/globularity of predicted structures for pLM-generated sequences – estimated as the fractional burial of total amino-acid surface area relative to the linear polypeptide chain – does not map onto whether generated variants are co-localizing with natural vs. random sequences (Fig. 3B). Similarly, SCOP fold labels are confidently assigned for large swaths of ProtGPT2-generated and ESM2-150M-generated sequences that do not co-localize with natural proteins (Fig. 3C). This suggests that both ProtGPT2 and ESM2-150M can emit sequences that are distinct in some statistical sense from natural ones yet able to fold into plausible and familiar 3D structures. This finding is tempered slightly by the realization that sequences from both PLMs are predicted to adopt structures that are less compact and less rich in secondary structure content (*α*-helix and *β*-sheet) on average than natural proteins in the SCOP reference set, implying that PLM output may tilt towards disordered regions (Fig. 3D-E). However, this latter effect may be exaggerated by the disparity between fixed generated sequence lengths and human-curated experimental structures with disordered head/tail regions that are not resolved and/or removed when assigning domain boundaries.

### PLM-generated structures do not follow the natural distribution

For a finer-grained perspective on structure, we look to the SCOP fold label assignment procedure and observe that while a respectable 32.7% of ProtGPT2-generated sequences are assignable to a fold label, this is only the case for 5.5% of ESM2-150M-generated sequences, on par with the 6.1% assignment rate for random sequences (Fig. 4A). That the ESM2-150M fold assignment rate is no improvement over a control approach that includes first-order sequence statistics sparks doubt as to whether Gibbs sampling can reflect the higher-order sequence correlations presumably learned by ESM2-150M without rejecting free generation in favor of a generation task that mimics the increased availability of contextual sequence information during the training task, which never drops below 15% of positions masked.

**Fig 4.**
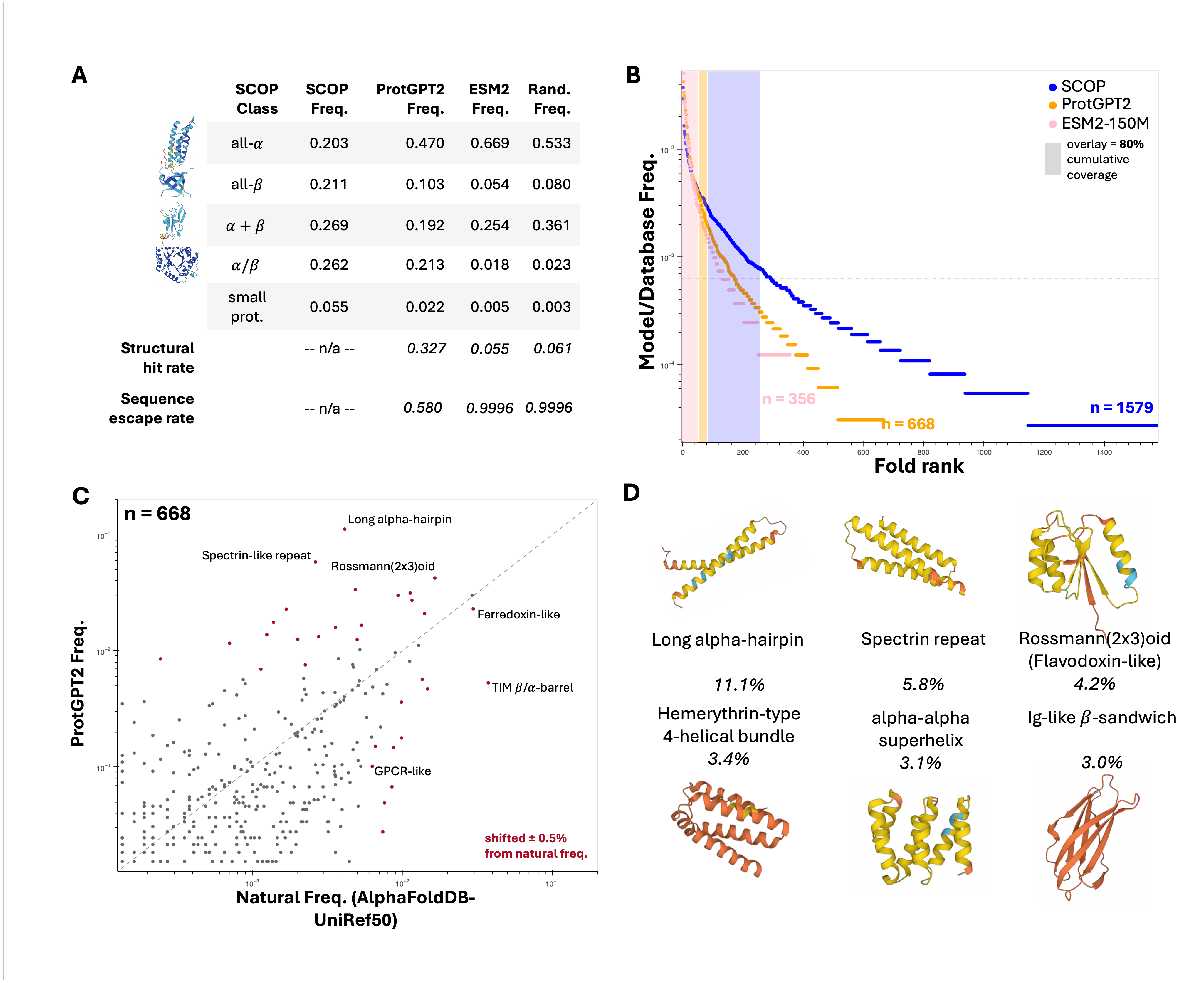
Structural ensembles generated by pretrained language models cover natural protein-space imperfectly. (**A**) Comparison of global protein topology preferences of natural, pLM-generated, and random sequences. (**B**)) Rank-ordered fold ensemble frequency plots for natural and pLM-generated sequences. (**C**) Fold ensemble comparison of ProtGPT2-generated sequences vs natural (SCOP-UnitRef50) sequences. **(D)** The 6 most-common SCOP folds among ProtGPT2 outputs; representative structures are of far-from-natural sequences (no MMseqs2 hit with E-value *<* 0.01).

Fold label assignments for both ESM2-150M and random sequences also skew heavily towards all-*α* topologies like helical bundles and *α* + *β* topologies like ferredoxins (Fig. 4A). SCOP topology class coverage with ProtGPT2 bears more resemblance to the natural distribution, especially as far as reaching the *α/β* folds that include most enzymatic diversity, but still overweights all-*α* content (Fig. 4A) [31]. These trends in structural coverage breadth propagate to the fold level; 668/1579 (42.3%) of SCOP folds are detected in ProtGPT2 output, or ∼ 1.9x the 356/1579 (22.5%) represented in ESM2-150M output (Fig. 4B). Focusing on ProtGPT2, overrepresented folds include several flavors of *α*-helical bundles, Rossmann(2×3)oids, and the all-*β* immunoglobulin-like domain, while underrepresented folds include ubiquitous and diversified functional folds such as TIM *β/α* barrels, and G-protein coupled receptors (GPCRs) (Fig. 4C-D, Table S.2). Evidently, plucked off the shelf, PLMs do not reproduce the natural frequencies of known protein folds.

### Prioritizing sequence novelty shrinks accessible structure-space

While the structural ensembles sampled by PLMs fail to cover the breadth of natural structural-space and distort frequencies in the corners that they do touch, the plausible structures that they *do* access come with notable sequence novelty. In particular, out of the 32,694/99,982 (32.7%) ProtGPT2-generated sequences with a fold label assignment, a further 18,962 (58.0% of assignable; 19.0% of all) have *no detectable homology* to any of the ∼ 50 million representative protein sequences in UniRef50, a phenomenon that we dub sequence “escape” (Fig. 4A, Table 1). One hypothesis, inspired by typical natural language processing (NLP) approaches, is that higher rates of sequence escape, and perhaps some of the missing structure coverage, might be reached by loosening sampling hyperparameters to encourage diversity in generated protein “text.” Continuing with ProtGPT2, the two critical and tunable hyperparameters are top_k and sampling temperature – increasing top_k allows for more tokens to be considered for sampling at a given step, while increasing temperature flattens the probability distribution over the token pool under consideration – both leading in theory to greater diversity in sequence output.

**Table 1.** Base ProtGPT2 sequence and structure generation performance depends on sampling hyperparameters. As sampling temperature and vocabulary size increase, generated sequences are more likely to lack homology to natural proteins, but also more likely to be unstructured and/or unassignable to any categorized SCOP fold label.

| Hyperparams |  | Results |  |  |  |
| --- | --- | --- | --- | --- | --- |
| top_k | temp | Valid Seq. | # Folds | Struct. Hit | Seq. Esc. |
| 600 | 0.800 | 1.000 | 658 | 0.347 | 0.445 |
|  | 1.000 | 1.000 | 635 | 0.336 | 0.545 |
|  | 1.200 | 1.000 | 645 | 0.322 | 0.629 |
|  | 1.500 | 1.000 | 617 | 0.304 | 0.717 |
|  | 2.000 | 0.999 | 606 | 0.282 | 0.797 |
|  | 5.000 | 0.981 | 513 | 0.160 | 0.912 |
| 950 | 0.800 | 1.000 | 643 | 0.345 | 0.466 |
|  | 1.000 | 1.000 | 668 | 0.327 | 0.580 |
|  | 1.200 | 0.999 | 620 | 0.306 | 0.674 |
|  | 1.500 | 1.000 | 625 | 0.287 | 0.766 |
|  | 2.000 | 0.998 | 587 | 0.262 | 0.855 |
|  | 5.000 | 0.985 | 473 | 0.151 | 0.958 |
| 1500 | 0.800 | 1.000 | 649 | 0.340 | 0.484 |
|  | 1.000 | 1.000 | 646 | 0.315 | 0.609 |
|  | 1.200 | 0.999 | 627 | 0.290 | 0.708 |
|  | 1.500 | 1.000 | 608 | 0.263 | 0.816 |
|  | 2.000 | 0.998 | 577 | 0.239 | 0.903 |
|  | 5.000 | 0.988 | 476 | 0.144 | 0.981 |
| 2400 | 0.800 | 1.000 | 634 | 0.334 | 0.493 |
|  | 1.000 | 1.000 | 634 | 0.303 | 0.628 |
|  | 1.200 | 1.000 | 617 | 0.277 | 0.742 |
|  | 1.500 | 1.000 | 588 | 0.248 | 0.857 |
|  | 2.000 | 0.998 | 542 | 0.222 | 0.944 |
|  | 5.000 | 0.991 | 460 | 0.139 | 0.993 |
| 4000 | 0.800 | 1.000 | 662 | 0.334 | 0.510 |
|  | 1.000 | 1.000 | 644 | 0.298 | 0.650 |
|  | 1.200 | 0.999 | 618 | 0.271 | 0.778 |
|  | 1.500 | 1.000 | 574 | 0.238 | 0.894 |
|  | 2.000 | 0.998 | 540 | 0.212 | 0.968 |
|  | 5.000 | 0.993 | 442 | 0.145 | 0.998 |

We systematically vary both temperature (*T* = 0.8, 1.0, 1.2, 1.5, 2.5, 5.0) and top_k (*N*_*k*_ = 600, 950, 1500, 2400, 4000), generating 100,000 sequences from ProtGPT2 for each of the 30 hyperparameter pairs on this grid and following the same truncation, filtering, structure prediction, and annotation workflow described previously. Consistent with the NLP hypothesis, we see that sequence escape rates increase dramatically when temperature or top_k is increased, and approach 100% of assignable structures when both are increased simultaneously; this trend holds in aggregate and at the level of individual fold classes (Table 1, Fig. 5). However, far from rescuing the missing structural breadth, boosting sequence novelty exacerbates the issue. As temperature and/or top_k are increased, the number of unique SCOP folds detected plummets, the fraction of structures assignable to any label (the “structural hit rate”) falls precipitously, and topology class representation vanishes in favor of all-*α* helical bundles, largely at the expense of *α/β* proteins (Table 1, Fig. S.1-S.2). Again, these trends propagate down to individual fold classes, with a handful of helical bundles dominating the generative space, albeit with impressive sequence escape rates (Tables S.1-S.5). While obtaining far-from-natural versions of helical bundles could yet prove useful for protein design writ large (e.g. in minibinder design campaigns), the structural biases accentuated by prioritizing sequence novelty reinforce the reality that without additional tuning or optimization of models and generative tasks, pretrained PLMs are at best flawed mirrors of natural protein-space thanks to severe structural dropout and a closely coupled tendency to fall into all-*α* topological traps.

**Fig 5.**
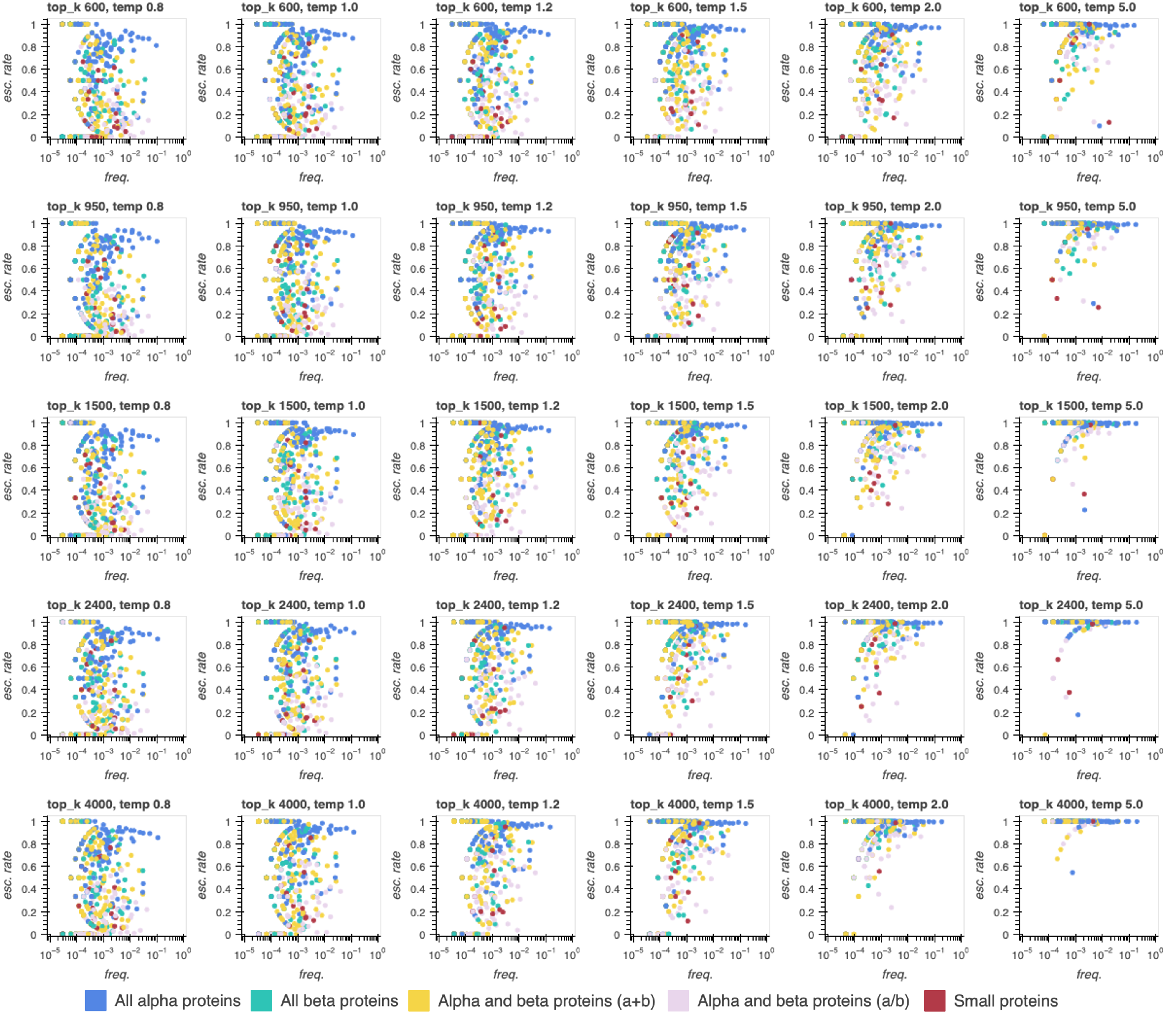
Sequence escape rates increase across most folds as sampling temperature increases, at the cost of a shift towards all-*α* topologies. Sequence escape rates for all assigned SCOP folds generated from ProtGPT2 within batches of 100k sequences for several sampling temperatures (0.8, 1, 1.2, 1.5, 2, 5) x several top_k values (number of highest-probability tokens considered in sampling out of 50,256 total; 600, 950, 1500, 2400, 4000.

## Discussion

We showed that, after “seeing” tens of millions of real protein sequences, PLMs are sufficiently aware of the sequence and structure statistics of natural proteins to yield realistic proteins that pass basic *in silico* biophysical screens – generated proteins are expected to be compact, globular, contain familiar secondary elements, and often bear resemblance to known structural motifs. This behavior emerges seamlessly for autoregressive models – exemplified by ProtGPT2 – where the piecewise left-to-right free sequence generation task echoes the training task; it is much more limited for ESM2-150M, a model trained with bidirectional partial masking and consequently likely better suited for descriptive and variant effect predictions than for sequence generation absent user-imposed constraints. We also demonstrated that ProtGPT2 is a powerful instrument for accessing sequence novelty, specifically sequences devoid of measurable similarity to natural proteins under highly sensitive homology search conditions.

However, this sequence novelty comes at a substantial cost – limited structural breadth in model output, sacrificing much of the order and richness of nature’s structural landscape. This presents as a fundamental tradeoff. The more sequence novelty is pursued by tuning sampling hyperparameters to explore the vastness of sequence-space, the more complete the collapse to a small collection of structural modes, largely biophysically simple *α*-helical bundles at that.

In contrast to situations encountered in foundational ML subfields including natural language processing and computer vision, this collapse to a subset of modes occurs *without* training data contamination and only weakly reflects the relative frequencies of these modes in the UniRef50 training data common to both ProtGPT2 and the ESM2 model family (versions 2021 04, and 2021 09, respectively) [32, 33]. Instead, this behavior may stem from a combination of limitations baked into model architecture (e.g. the unidirectional context window of ProtGPT2) and mechanistic discordance between training and generation tasks (e.g. 15% vs 100% sequence masking in training vs. generation contexts respectively for ESM2). The overrepresentation of a limited pool of structural motifs and the concordant absence of many rare, functionally relevant, and/or evolvable ones from generative PLM output could prove deleterious for future AI-driven protein design campaigns. Alternative forcing strategies will be required for reliably sampling functional-yet-novel protein populations from PLMs, particularly in the pursuit of “linguistically consistent” proteins well beyond the confines of natural sequence-space, an essential goal for high-throughput design of related sets of bespoke ligands and biocatalysts.

## Materials and methods

Except where otherwise specified, all model access and interfacing was via trill v1.3.11 [34].

### Sequence Generation from Protein Language Models

For the model comparison experiment, sequences (*n* = 100, 000) were sampled from ProtGPT2 by L-to-R next-token prediction with the default best-performing hyperparameters from ref. [2]; sampling temperature 1, top_k 950, top p 1.0, repetition penalty 1.2. The termination condition was set following the 40th token or the first STOP token occurring prior to the 40th token; sequences longer than 100aa were truncated to 100aa as the maximal length. Sequences containing rare or ambiguous amino acids (B, J, O, U, X, or Z) were filtered out as invalid, leaving 99,982 sequences. Sequences were sampled from ESM2-150M (*n* = 148, 500), from L-to-R with next-token prediction with Gibbs sampling, with a default sampling temperature of 1, no repetition penalty, and allowing for sampling from the full token distribution. The termination condition was set following the 100th amino-acid or the first STOP token occurring prior to the 100th amino-acid. Following deduplication and truncation and filtering applied as for ProtGPT2, 144,043 sequences were carried through for downstream analysis.

For the hyperparameter scan experiment, sequences (*n* = 100, 000 per configuration) were generated from ProtGPT2 by L-to-R next-token prediction with top p 1.0 and repetition penalty 1.2 fixed, and a grid search over 30 (temperature, top_k) pairs derived from six possible temperatures (*T* = 0.8, 1.0, 1.2, 1.5, 2.0, 5.0) x five possible top_k pool sizes (*N*_*k*_ = 600, 950, 1500, 2400, 4000). Truncation and filtering were applied as in the model comparison experiment.

### Sequence Generation from Control Models

The random-sequence control set was generated by position-independent sampling of *n* = 74, 250 sequences of length 100aa from the 20 proteinogenic amino acids, with sampling probability for each amino acid proportional to its natural abundance. As sequence length was fixed and the rare/ambiguous amino acids B, J, O, U, X, and Z excluded, no filtering or truncation steps were required.

The inverse-folding control set was constructed by generating three sequences from ESM-IF1 with each of the 36,900 representative structures in the SCOP database as a backbone template, for *n* = 110, 700 sequences in total. Pre- and post-processing for rare ligands in templates and rare/ambiguous amino acids in outputs, respectively, reduced inverse-folding output to 104,591 sequences. Default hyperparameters for sampling were taken as in ref. [24].

### Validation of ESMFold on Out-of-Distribution Samples

A validation set for ESMFold structure prediction accuracy on out-of-distribution sequences with respect to natural UniRef50 training data was assembled from *de novo* proteins (“artificial sequence” taxonomy) with structures deposited in the Protein Data Bank (PDB) on-or-after the ESMFold training cutoff date of 05-01-2020 and before 11-01-2023. Mirroring the training set construction process described in [1], we filtered out structures with resolution > 9 Å, length ≤ 20aa, rare or ambiguous amino acids (BJOUXZ), or containing > 20% sequence composition of any one amino acid, and clustered remaining sequences at the 40% identity level to obtain the final validation set of *n* = 122 sequences.

### Structure Prediction and Assignment

All structures (for filtered, truncated sequences as described above) were predicted with default ESMFold inference parameters as in ref/ [1]. For the model comparison experiment, structures were singly-inferenced (batch size 1), with compute resource collaboration with Yurts AI (now Legion Intelligence). For the hyperparameter scan experiment, structures were batch-inferenced with batch size 100 to optimally utilize memory allocation on A100-80GB GPUs, with compute resource collaboration through Oracle Cloud Infrastructure (OCI).

Predicted structures were annotated to SCOP fold labels via Foldseek structure-based search (running in accelerated TMalign mode) against a reference database containing the *n* = 36, 900 SCOP superfamily representative domains retrieved from the PDB. The consensus SCOP fold was defined as the fold accounting for the most hits with TMscore > 0.5 and max(query coverage, target coverage) > 0.8.

### Sensitive Sequence Search and Novelty Characterization

In both the model comparison and hyperparameter scan experiments, PLM-generated and control sequences were searched against UniRef50 using mmseqs2 with default easy-search parameters and maximum e-value 0.01. Sequence escape rate was computed as the fraction of sequences not returning an alignment hit of any length to any cluster representative from UniRef50 at the specified e-value threshold.

### Construction of the SCOP-UniRef50 Sequence-Structure Database

The SCOP-UniRef50 custom sequence-structure fragment database was constructed by performing reciprocal Foldseek searches (in fast TM-align mode) of the SCOP database of superfamily representative PDB structures (*n* = 36, 900) against the UniRef50 portion (based on the 2021 04 release) included in the July 2022 update to the AlphaFoldDB as first reported in [28] and made available as a precompiled Foldseek database in [27], filtering for reciprocal hits with fractional query and target coverage > 0.8 and TMscore> 0.5, and clustering the filtered fragments at 100% identity.

For the model comparison experiments, *n* = 100, 000 natural sequences were uniformly sampled from SCOP-UniRef50 and jointly embedded along with PLM-generated and control sequences using ESM2-150M. This choice was made vs. sampling directly from SCOP in order to: (1) obtain a similar number of natural sequences (∼ 10^5^) to model-generated and control batches; (2) draw sequence fragments with representative taxonomic coverage for evolutionarily conserved folds, as opposed to the narrower taxonomic coverage in SCOP, itself a function of skewed taxonomic coverage in the Protein Data Bank [25].

### Basic Chemical Property Calculations

Amino-acid surface area burial fraction was calculated using custom code and reference individual amino-acid surface areas (HMS Bionumbers: 103239). Secondary structure annotations were assigned with DSSP via the corresponding PyMOL v3.1.0 wrapper.

## Supporting information

### Supplemental Figures

**Fig S.1.**
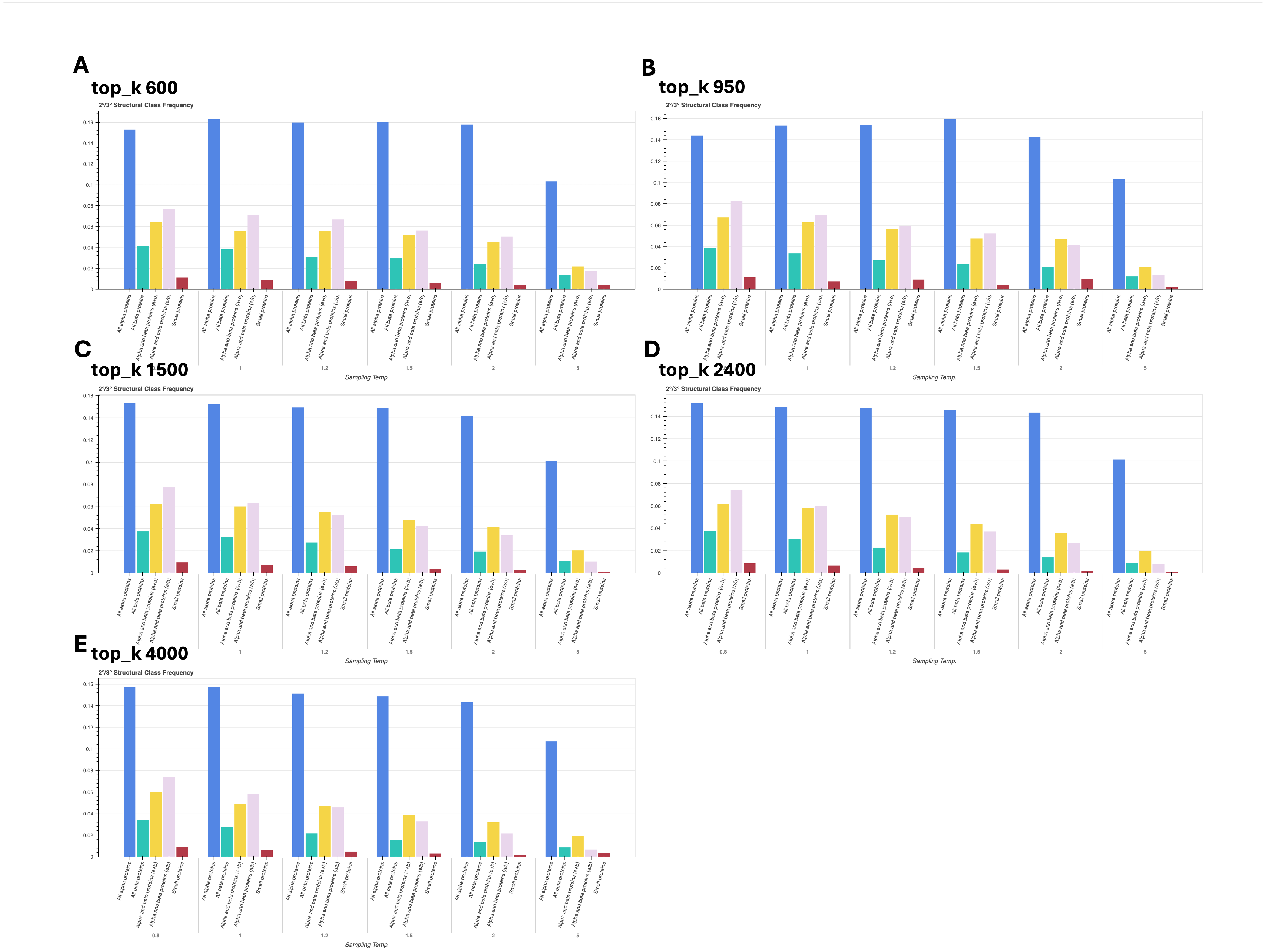
Structure hit rates from base ProtGPT2 decrease as sampling temperature and top_k increase. Structure hit rates from batches of 100k sequences generated from ProtGPT2 for several sampling temperatures (0.8, 1, 1.2, 1.5, 2, 5) and top_k values (number of highest-probability tokens considered in sampling out of 50,256 total) – (**A**) 600, (**B**) 950, (**C**) 1500, (**D**) 2400, (**E**) 4000; broken down by protein global topology class (*α, β, α* + *β, α/β*, or “small / minimal 2° structure”)

**Fig S.2.**
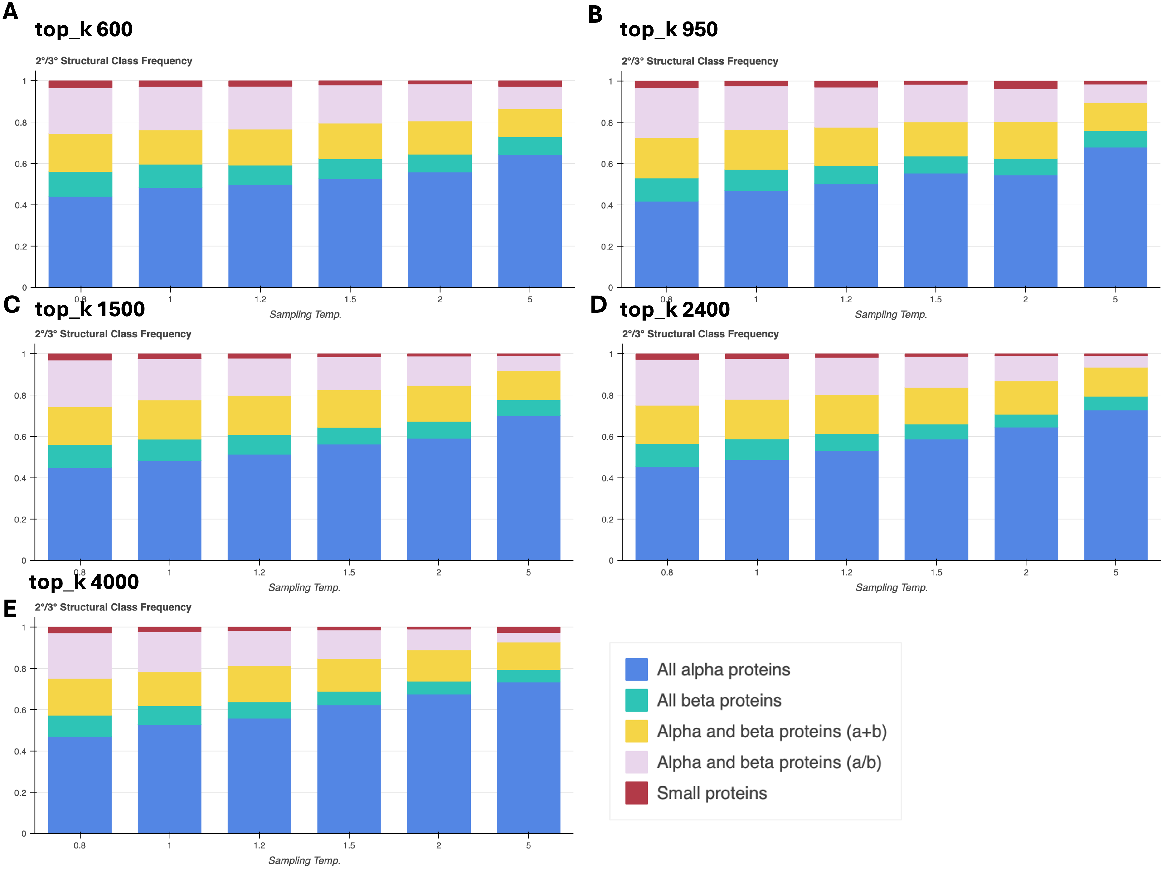
Generated fold distributions shift towards all-*α* proteins and away from *α/β* proteins as sampling temperature increases. Frequency of each protein global topology class (*α, β, α* + *β, α/β*, or “small / minimal 2° structure”) among all structure hits within batches of 100k sequences generated from ProtGPT2 for several sampling temperatures (0.8, 1, 1.2, 1.5, 2, 5) and top_k values (number of highest-probability tokens considered in sampling out of 50,256 total) – (**A**) 600, (**B**) 950, (**C**) 1500, (**D**) 2400, (**E**) 4000

### Supplemental Tables

**Table S.1.** Most common SCOP folds generated by base ProtGPT2 at various sampling temperatures with top_k 600.

| temp: 0.8 |  |  |  |  |
| --- | --- | --- | --- | --- |
| Fold | Class | Freq. | Abs. Hit Rate | Esc. Rate |
| Long alpha-hairpin | $\alpha$ | 0.095 | 0.033 | 0.810 |
| Spectrin repeat-like | $\alpha$ | 0.050 | 0.017 | 0.871 |
| Rossmann(2x3)oid (Flavodoxin-like) | $\alpha/\beta$ | 0.048 | 0.017 | 0.146 |
| Immunoglobulin-like beta-sandwich | $\beta$ | 0.036 | 0.012 | 0.510 |
| alpha-alpha superhelix | $\alpha$ | 0.033 | 0.011 | 0.386 |
| temp: 1 |  |  |  |  |
| Fold | Class | Freq. | Abs. Hit Rate | Esc. Rate |
| Long alpha-hairpin | $\alpha$ | 0.112 | 0.038 | 0.874 |
| Spectrin repeat-like | $\alpha$ | 0.056 | 0.019 | 0.907 |
| Rossmann(2x3)oid (Flavodoxin-like) | $\alpha/\beta$ | 0.043 | 0.014 | 0.252 |
| Immunoglobulin-like beta-sandwich | $\beta$ | 0.034 | 0.012 | 0.596 |
| Hemerythrin-type up-and-down 4-helical bundle | $\alpha$ | 0.032 | 0.011 | 0.903 |
| temp: 1.2 |  |  |  |  |
| Fold | Class | Freq. | Abs. Hit Rate | Esc. Rate |
| Long alpha-hairpin | $\alpha$ | 0.122 | 0.039 | 0.908 |
| Spectrin repeat-like | $\alpha$ | 0.063 | 0.020 | 0.929 |
| Rossmann(2x3)oid (Flavodoxin-like) | $\alpha/\beta$ | 0.043 | 0.014 | 0.330 |
| Hemerythrin-type up-and-down 4-helical bundle | $\alpha$ | 0.036 | 0.012 | 0.937 |
| Immunoglobulin-like beta-sandwich | $\beta$ | 0.030 | 0.010 | 0.696 |
| temp: 1.5 |  |  |  |  |
| Fold | Class | Freq. | Abs. Hit Rate | Esc. Rate |
| Long alpha-hairpin | $\alpha$ | 0.130 | 0.040 | 0.943 |
| Spectrin repeat-like | $\alpha$ | 0.068 | 0.021 | 0.950 |
| Rossmann(2x3)oid (Flavodoxin-like) | $\alpha/\beta$ | 0.042 | 0.013 | 0.424 |
| Hemerythrin-type up-and-down 4-helical bundle | $\alpha$ | 0.040 | 0.012 | 0.958 |
| Immunoglobulin/albumin-binding domain-like | $\alpha$ | 0.033 | 0.010 | 0.955 |
| temp: 2 |  |  |  |  |
| Fold | Class | Freq. | Abs. Hit Rate | Esc. Rate |
| Long alpha-hairpin | $\alpha$ | 0.141 | 0.040 | 0.969 |
| Spectrin repeat-like | $\alpha$ | 0.075 | 0.021 | 0.968 |
| Hemerythrin-type up-and-down 4-helical bundle | $\alpha$ | 0.044 | 0.013 | 0.978 |
| Rossmann(2x3)oid (Flavodoxin-like) | $\alpha/\beta$ | 0.043 | 0.012 | 0.500 |
| Immunoglobulin/albumin-binding domain-like | $\alpha$ | 0.032 | 0.009 | 0.966 |
| temp: 5 |  |  |  |  |
| Fold | Class | Freq. | Abs. Hit Rate | Esc. Rate |
| Long alpha-hairpin | $\alpha$ | 0.154 | 0.025 | 0.989 |
| Spectrin repeat-like | $\alpha$ | 0.084 | 0.014 | 0.989 |
| Hemerythrin-type up-and-down 4-helical bundle | $\alpha$ | 0.060 | 0.010 | 0.984 |
| alpha-alpha superhelix | $\alpha$ | 0.040 | 0.006 | 0.905 |
| Immunoglobulin/albumin-binding domain-like | $\alpha$ | 0.033 | 0.005 | 0.987 |

**Table S.2.** Most common SCOP folds generated by base ProtGPT2 at various sampling temperatures with top_k (vocabulary size) 950.

| temp: 0.8 |  |  |  |  |
| --- | --- | --- | --- | --- |
| Fold | Class | Freq. | Abs. Hit Rate | Esc. Rate |
| Long alpha-hairpin | $\alpha$ | 0.101 | 0.035 | 0.841 |
| Spectrin repeat-like | $\alpha$ | 0.050 | 0.017 | 0.872 |
| Rossmann(2x3)oid (Flavodoxin-like) | $\alpha/\beta$ | 0.048 | 0.016 | 0.177 |
| Immunoglobulin-like beta-sandwich | $\beta$ | 0.032 | 0.011 | 0.534 |
| Canonical WHD (winged helix domain) fold | $\alpha + \beta$ | 0.031 | 0.011 | 0.342 |
| temp: 1 |  |  |  |  |
| Fold | Class | Freq. | Abs. Hit Rate | Esc. Rate |
| Long alpha-hairpin | $\alpha$ | 0.111 | 0.036 | 0.893 |
| Spectrin repeat-like | $\alpha$ | 0.058 | 0.019 | 0.918 |
| Rossmann(2x3)oid (Flavodoxin-like) | $\alpha/\beta$ | 0.042 | 0.014 | 0.273 |
| Hemerythrin-type up-and-down 4-helical bundle | $\alpha$ | 0.034 | 0.011 | 0.926 |
| alpha-alpha superhelix | $\alpha$ | 0.031 | 0.010 | 0.571 |
| temp: 1.2 |  |  |  |  |
| Fold | Class | Freq. | Abs. Hit Rate | Esc. Rate |
| Long alpha-hairpin | $\alpha$ | 0.126 | 0.039 | 0.930 |
| Spectrin repeat-like | $\alpha$ | 0.065 | 0.020 | 0.946 |
| Rossmann(2x3)oid (Flavodoxin-like) | $\alpha/\beta$ | 0.041 | 0.013 | 0.345 |
| Hemerythrin-type up-and-down 4-helical bundle | $\alpha$ | 0.038 | 0.012 | 0.948 |
| Canonical WHD (winged helix domain) fold | $\alpha + \beta$ | 0.030 | 0.009 | 0.530 |
| temp: 1.5 |  |  |  |  |
| Fold | Class | Freq. | Abs. Hit Rate | Esc. Rate |
| Long alpha-hairpin | $\alpha$ | 0.136 | 0.039 | 0.943 |
| Spectrin repeat-like | $\alpha$ | 0.069 | 0.020 | 0.969 |
| Hemerythrin-type up-and-down 4-helical bundle | $\alpha$ | 0.046 | 0.013 | 0.960 |
| Rossmann(2x3)oid (Flavodoxin-like) | $\alpha/\beta$ | 0.042 | 0.012 | 0.491 |
| Immunoglobulin/albumin-binding domain-like | $\alpha$ | 0.030 | 0.009 | 0.960 |
| temp: 2 |  |  |  |  |
| Fold | Class | Freq. | Abs. Hit Rate | Esc. Rate |
| Long alpha-hairpin | $\alpha$ | 0.149 | 0.039 | 0.978 |
| Spectrin repeat-like | $\alpha$ | 0.076 | 0.020 | 0.984 |
| Hemerythrin-type up-and-down 4-helical bundle | $\alpha$ | 0.045 | 0.012 | 0.976 |
| Rossmann(2x3)oid (Flavodoxin-like) | $\alpha/\beta$ | 0.040 | 0.010 | 0.596 |
| Immunoglobulin/albumin-binding domain-like | $\alpha$ | 0.035 | 0.009 | 0.974 |
| temp: 5 |  |  |  |  |
| Fold | Class | Freq. | Abs. Hit Rate | Esc. Rate |
| Long alpha-hairpin | $\alpha$ | 0.178 | 0.027 | 0.991 |
| Spectrin repeat-like | $\alpha$ | 0.090 | 0.014 | 0.996 |
| Hemerythrin-type up-and-down 4-helical bundle | $\alpha$ | 0.064 | 0.010 | 0.989 |
| Immunoglobulin/albumin-binding domain-like | $\alpha$ | 0.038 | 0.006 | 0.986 |
| alpha-alpha superhelix | $\alpha$ | 0.035 | 0.005 | 0.934 |

**Table S.3.** Most common SCOP folds generated by base ProtGPT2 at various sampling temperatures with top_k (vocabulary size) 1500.

| temp: 0.8 |  |  |  |  |  |
| --- | --- | --- | --- | --- | --- |
| Fold | Class | Freq. | Abs. Hit Rate | Esc. Rate |  |
| Long alpha-hairpin | $\alpha$ | 0.101 | 0.035 | 0.847 | |
| Spectrin repeat-like | $\alpha$ | 0.052 | 0.018 | 0.878 | |
| Rossmann(2x3)oid (Flavodoxin-like) | $\alpha/\beta$ | 0.045 | 0.015 | 0.210 | |
| Immunoglobulin-like beta-sandwich | $\beta$ | 0.033 | 0.011 | 0.555 | |
| alpha-alpha superhelix | $\alpha$ | 0.031 | 0.010 | 0.426 | |
| temp: 1 |  |  |  |  |  |
| Fold | Class | Freq. | Abs. Hit Rate | Esc. Rate |  |
| Long alpha-hairpin | $\alpha$ | 0.119 | 0.037 | 0.895 | |
| Spectrin repeat-like | $\alpha$ | 0.059 | 0.019 | 0.918 | |
| Rossmann(2x3)oid (Flavodoxin-like) | $\alpha/\beta$ | 0.040 | 0.013 | 0.304 | |
| Hemerythrin-type up-and-down 4-helical bundle | $\alpha$ | 0.035 | 0.011 | 0.930 | |
| Canonical WHD (winged helix domain) fold | $\alpha + \beta$ | 0.029 | 0.009 | 0.472 | |
| temp: 1.2 |  |  |  |  |  |
| Fold | Class | Freq. | Abs. Hit Rate | Esc. Rate |  |
| Long alpha-hairpin | $\alpha$ | 0.129 | 0.038 | 0.930 | |
| Spectrin repeat-like | $\alpha$ | 0.067 | 0.019 | 0.956 | |
| Rossmann(2x3)oid (Flavodoxin-like) | $\alpha/\beta$ | 0.042 | 0.012 | 0.425 | |
| Hemerythrin-type up-and-down 4-helical bundle | $\alpha$ | 0.040 | 0.012 | 0.951 | |
| Canonical WHD (winged helix domain) fold | $\alpha + \beta$ | 0.029 | 0.008 | 0.528 | |
| temp: 1.5 |  |  |  |  |  |
| Fold | Class | Freq. | Abs. Hit Rate | Esc. Rate |  |
| Long alpha-hairpin | $\alpha$ | 0.145 | 0.038 | 0.963 | |
| Spectrin repeat-like | $\alpha$ | 0.077 | 0.020 | 0.984 | |
| Hemerythrin-type up-and-down 4-helical bundle | $\alpha$ | 0.047 | 0.012 | 0.976 | |
| Rossmann(2x3)oid (Flavodoxin-like) | $\alpha/\beta$ | 0.040 | 0.011 | 0.566 | |
| Immunoglobulin/albumin-binding domain-like | $\alpha$ | 0.033 | 0.009 | 0.968 | |
| temp: 2 |  |  |  |  |  |
| Fold | Class | Freq. | Abs. Hit Rate | Esc. Rate |  |
| Long alpha-hairpin | $\alpha$ | 0.158 | 0.038 | 0.988 | |
| Spectrin repeat-like | $\alpha$ | 0.078 | 0.019 | 0.989 | |
| Hemerythrin-type up-and-down 4-helical bundle | $\alpha$ | 0.050 | 0.012 | 0.986 | |
| Rossmann(2x3)oid (Flavodoxin-like) | $\alpha/\beta$ | 0.039 | 0.009 | 0.708 | |
| Immunoglobulin/albumin-binding domain-like | $\alpha$ | 0.034 | 0.008 | 0.987 | |
| temp: 5 |  |  |  |  |  |
| Fold | Class | Freq. | Abs. Hit Rate | Esc. Rate |  |
| Long alpha-hairpin | $\alpha$ | 0.182 | 0.026 | 0.993 | |
| Spectrin repeat-like | $\alpha$ | 0.094 | 0.014 | 0.999 | |
| Hemerythrin-type up-and-down 4-helical bundle | $\alpha$ | 0.070 | 0.010 | 0.994 | |
| Ferredoxin-like | $\alpha + \beta$ | 0.040 | 0.006 | 0.984 | |
| Immunoglobulin/albumin-binding domain-like | $\alpha$ | 0.038 | 0.005 | 0.993 | |

**Table S.4.** Most common SCOP folds generated by base ProtGPT2 at various sampling temperatures with top_k (vocabulary size) 2400.

| temp: 0.8 |  |  |  |  |  |
| --- | --- | --- | --- | --- | --- |
| Fold | Class | Freq. | Abs. Hit Rate | Esc. Rate |  |
| Long alpha-hairpin | $\alpha$ | 0.106 | 0.036 | 0.850 | |
| Spectrin repeat-like | $\alpha$ | 0.052 | 0.018 | 0.894 | |
| Rossmann(2x3)oid (Flavodoxin-like) | $\alpha/\beta$ | 0.043 | 0.014 | 0.210 | |
| alpha-alpha superhelix | $\alpha$ | 0.032 | 0.011 | 0.435 | |
| Canonical WHD (winged helix domain) fold | $\alpha + \beta$ | 0.031 | 0.011 | 0.358 | |
| temp: 1 |  |  |  |  |  |
| Fold | Class | Freq. | Abs. Hit Rate | Esc. Rate |  |
| Long alpha-hairpin | $\alpha$ | 0.125 | 0.038 | 0.905 | |
| Spectrin repeat-like | $\alpha$ | 0.062 | 0.019 | 0.939 | |
| Rossmann(2x3)oid (Flavodoxin-like) | $\alpha/\beta$ | 0.040 | 0.012 | 0.353 | |
| Hemerythrin-type up-and-down 4-helical bundle | $\alpha$ | 0.038 | 0.011 | 0.917 | |
| Canonical WHD (winged helix domain) fold | $\alpha + \beta$ | 0.028 | 0.008 | 0.446 | |
| temp: 1.2 |  |  |  |  |  |
| Fold | Class | Freq. | Abs. Hit Rate | Esc. Rate |  |
| Long alpha-hairpin | $\alpha$ | 0.138 | 0.038 | 0.945 | |
| Spectrin repeat-like | $\alpha$ | 0.071 | 0.020 | 0.959 | |
| Hemerythrin-type up-and-down 4-helical bundle | $\alpha$ | 0.043 | 0.012 | 0.957 | |
| Rossmann(2x3)oid (Flavodoxin-like) | $\alpha/\beta$ | 0.041 | 0.011 | 0.456 | |
| Ferredoxin-like | $\alpha + \beta$ | 0.030 | 0.008 | 0.792 | |
| temp: 1.5 |  |  |  |  |  |
| Fold | Class | Freq. | Abs. Hit Rate | Esc. Rate |  |
| Long alpha-hairpin | $\alpha$ | 0.158 | 0.039 | 0.976 | |
| Spectrin repeat-like | $\alpha$ | 0.077 | 0.019 | 0.985 | |
| Hemerythrin-type up-and-down 4-helical bundle | $\alpha$ | 0.052 | 0.013 | 0.981 | |
| Rossmann(2x3)oid (Flavodoxin-like) | $\alpha/\beta$ | 0.038 | 0.010 | 0.601 | |
| Ferredoxin-like | $\alpha + \beta$ | 0.033 | 0.008 | 0.888 | |
| temp: 2 |  |  |  |  |  |
| Fold | Class | Freq. | Abs. Hit Rate | Esc. Rate |  |
| Long alpha-hairpin | $\alpha$ | 0.167 | 0.037 | 0.994 | |
| Spectrin repeat-like | $\alpha$ | 0.086 | 0.019 | 0.992 | |
| Hemerythrin-type up-and-down 4-helical bundle | $\alpha$ | 0.056 | 0.012 | 0.991 | |
| Immunoglobulin/albumin-binding domain-like | $\alpha$ | 0.041 | 0.009 | 0.991 | |
| Ferredoxin-like | $\alpha + \beta$ | 0.036 | 0.008 | 0.956 | |
| temp: 5 |  |  |  |  |  |
| Fold | Class | Freq. | Abs. Hit Rate | Esc. Rate |  |
| Long alpha-hairpin | $\alpha$ | 0.190 | 0.027 | 0.998 | |
| Spectrin repeat-like | $\alpha$ | 0.095 | 0.013 | 0.998 | |
| Hemerythrin-type up-and-down 4-helical bundle | $\alpha$ | 0.069 | 0.010 | 0.998 | |
| Ferredoxin-like | $\alpha + \beta$ | 0.041 | 0.006 | 0.993 | |
| Immunoglobulin/albumin-binding domain-like | $\alpha$ | 0.041 | 0.006 | 0.996 | |

**Table S.5.** Most common SCOP folds generated by base ProtGPT2 at various sampling temperatures with top_k (vocabulary size) 4000.

| temp: 0.8 |  |  |  |  |
| --- | --- | --- | --- | --- |
| Fold | Class | Freq. | Abs. Hit Rate | Esc. Rate |
| Long alpha-hairpin | $\alpha$ | 0.108 | 0.036 | 0.855 |
| Spectrin repeat-like | $\alpha$ | 0.054 | 0.018 | 0.892 |
| Rossmann(2x3)oid (Flavodoxin-like) | $\alpha/\beta$ | 0.042 | 0.014 | 0.217 |
| alpha-alpha superhelix | $\alpha$ | 0.031 | 0.010 | 0.448 |
| Hemerythrin-type up-and-down 4-helical bundle | $\alpha$ | 0.031 | 0.010 | 0.896 |
| temp: 1 |  |  |  |  |
| Fold | Class | Freq. | Abs. Hit Rate | Esc. Rate |
| Long alpha-hairpin | $\alpha$ | 0.123 | 0.037 | 0.904 |
| Spectrin repeat-like | $\alpha$ | 0.065 | 0.019 | 0.939 |
| Rossmann(2x3)oid (Flavodoxin-like) | $\alpha/\beta$ | 0.039 | 0.012 | 0.377 |
| Hemerythrin-type up-and-down 4-helical bundle | $\alpha$ | 0.038 | 0.011 | 0.930 |
| alpha-alpha superhelix | $\alpha$ | 0.028 | 0.008 | 0.609 |
| temp: 1.2 |  |  |  |  |
| Fold | Class | Freq. | Abs. Hit Rate | Esc. Rate |
| Long alpha-hairpin | $\alpha$ | 0.146 | 0.039 | 0.949 |
| Spectrin repeat-like | $\alpha$ | 0.071 | 0.019 | 0.974 |
| Hemerythrin-type up-and-down 4-helical bundle | $\alpha$ | 0.046 | 0.012 | 0.967 |
| Rossmann(2x3)oid (Flavodoxin-like) | $\alpha/\beta$ | 0.041 | 0.011 | 0.544 |
| Ferredoxin-like | $\alpha + \beta$ | 0.031 | 0.008 | 0.812 |
| temp: 1.5 |  |  |  |  |
| Fold | Class | Freq. | Abs. Hit Rate | Esc. Rate |
| Long alpha-hairpin | $\alpha$ | 0.161 | 0.038 | 0.981 |
| Spectrin repeat-like | $\alpha$ | 0.086 | 0.020 | 0.991 |
| Hemerythrin-type up-and-down 4-helical bundle | $\alpha$ | 0.054 | 0.013 | 0.982 |
| Immunoglobulin/albumin-binding domain-like | $\alpha$ | 0.039 | 0.009 | 0.983 |
| Rossmann(2x3)oid (Flavodoxin-like) | $\alpha/\beta$ | 0.035 | 0.008 | 0.699 |
| temp: 2 |  |  |  |  |
| Fold | Class | Freq. | Abs. Hit Rate | Esc. Rate |
| Long alpha-hairpin | $\alpha$ | 0.183 | 0.039 | 0.997 |
| Spectrin repeat-like | $\alpha$ | 0.092 | 0.019 | 0.994 |
| Hemerythrin-type up-and-down 4-helical bundle | $\alpha$ | 0.062 | 0.013 | 0.998 |
| Immunoglobulin/albumin-binding domain-like | $\alpha$ | 0.038 | 0.008 | 0.995 |
| Ferredoxin-like | $\alpha + \beta$ | 0.038 | 0.008 | 0.970 |
| temp: 5 |  |  |  |  |
| Fold | Class | Freq. | Abs. Hit Rate | Esc. Rate |
| Long alpha-hairpin | $\alpha$ | 0.196 | 0.029 | 0.999 |
| Spectrin repeat-like | $\alpha$ | 0.097 | 0.014 | 1.000 |
| Hemerythrin-type up-and-down 4-helical bundle | $\alpha$ | 0.079 | 0.011 | 1.000 |
| Ferredoxin-like | $\alpha + \beta$ | 0.040 | 0.006 | 0.998 |
| Immunoglobulin/albumin-binding domain-like | $\alpha$ | 0.038 | 0.005 | 1.000 |

## Acknowledgments

We thank Steve Mayo, Erik Winfree, Richard Murray, Alec Lourenço, Andrew DeLaitsch, Mikel Lipschitz, and William Rosencrans, as well as all members of the Thomson Lab for thoughtful discussions and feedback.

This work was supported by the National Institutes of Health under award number R01GM150125, the Gordon and Betty Moore Foundation, the David & Lucille Packard Foundation under a Packard Fellowship to M.T., and the Heritage Medical Research Institute. The authors have no competing interests to disclose.

